# Swedish APP follows distinct neuronal trafficking itineraries that underlie its increased β-cleavage

**DOI:** 10.64898/2026.08.22.746059

**Authors:** Cameron R. Plowinske, Sarah Tedesco, Alec T. Nabb, Geraldine B. Quinones, Marvin Bentley

## Abstract

The amyloid precursor protein (APP) cleavage product Aβ comprises amyloid plaques in Alzheimer’s disease (AD). Aβ production is thought to occur in neuronal endosomes. Swedish APP (APP^Swe^) is associated with increased Aβ production and early onset AD, but it is unclear if APP^wt^ and APP^Swe^ differ in their neuronal trafficking. We performed quantitative live-cell microscopy with novel imaging-based assays in cultured hippocampal neurons to determine APP trafficking pathways. APP^wt^ and APP^Swe^ differed in their trafficking. APP^Swe^ was sorted into an additional vesicle population at the trans-Golgi. APP^Swe^ that reached the dendritic plasma membrane was less likely to be targeted to lysosomes and more likely to transcytose to the axon. Finally, we determined that amyloidogenic cleavage of APP was not limited to endosomes but also occurred in Golgi-derived vesicles. These results indicate that motifs in the APP ectodomain direct its sorting and that increased Aβ production of APP^Swe^ is facilitated by its specific trafficking.

## INTRODUCTION

Proteolytic cleavage of amyloid precursor protein (APP) by β -site APP cleaving enzyme-1 (BACE-1) and λ - secretase generates amyloid-β (Aβ), which aggregates to form amyloid plaques in the brain and is a pathological hallmark of Alzheimer’s disease (AD) (O’Brien and Wong, 2011; Selkoe and Hardy, 2016). The only current treatment for AD involves slowing cognitive decline in patients by clearing Aβ plaques, showing that Aβ plays a causative role in AD. Specific mutations in APP are associated with early-onset AD and increased Aβ production. The most prominent example is Swedish APP (APP^Swe^) which has two point mutations (KM670/671NL) in its ectodomain, near the BACE-1 cleavage site (Citron et al., 1992; Haass et al., 1995). However, the specific trafficking pathways of APP in neurons, and if differences in the trafficking of APP^wt^ and APP^Swe^ contribute to increased Aβ production of APP^Swe^ are unknown.

Due to the importance of Aβ in AD, much work has focused on APP processing by proteases while less attention has been paid to its neuronal trafficking. Furthermore, nearly all relevant experiments were performed in non-neuronal cells, neuron-like cell lines, or iPSC-derived neurons that may not fully reconstitute the complex trafficking physiology of primary mammalian neurons, which are extreme in their polarity, geometry, and size (Bentley and Banker, 2016). In addition, APP has been identifiied in nearly all compartments of the endomembrane system, but the trafficking itinerary of APP—especially in neurons— remains poorly defiined (Sun and Roy, 2018; Goldstein and Das, 2021).

Because BACE-1-mediated APP cleavage determines APP processing into Aβ (Thinakaran and Koo, 2008), there has been effort to defiine the compartment in which BACE-1 cleaves APP. Three basic models have emerged: 1) APP is cleaved in the secretory pathway (Wang et al., 2024; Fourriere and Gleeson, 2021; Thinakaran et al., 1996), 2) APP is cleaved in transcytotic vesicles that move endocytosed APP from the dendritic membrane into the axon (Woodruff et al., 2016; Niederst et al., 2015), and 3) APP is cleaved in dendritic endosomes (Das et al., 2013, 2015). Because the experiments supporting each model were performed in different model systems—only the third being supported with data from primary neurons— integrating the existing data has been challenging. Furthermore, these models result from a range of methods, including biochemical assays and imaging of fixed specimens. These techniques cannot capture the complexity of trafficking in individual live cells. Membrane trafficking is an inherently dynamic process, and live-cell imaging is necessary to understand protein trafficking, particularly for proteins with complex trafficking behaviors, such as APP (Haass et al., 2012; Tan and Gleeson, 2019a).

In this study, we systematically compared endosomal targeting of APP between unpolarized cells and primary hippocampal neurons. We determined that APP targeting to endosomes was less prevalent in neurons. Transport analyses revealed that APP^Swe^ was sorted less stringently at the trans-Golgi and entered additional post-Golgi vesicles that were inaccessible to APP^wt^. In addition, we found differences in the sorting of APP^wt^ and APP^Swe^ after dendritic endocytosis. APP^wt^ was targeted to lysosomes more efficiently, whereas a higher proportion of APP^Swe^ bypassed lysosomes and underwent transcytosis to axons. These experiments show that trafficking differences between APP^wt^ and APP^Swe^ likely contribute to the increase in Aβ generation that is associated with APP^Swe^ and that this trafficking difference is specific to neurons.

## RESULTS

### APP localization in neurons differs substantially from its localization in unpolarized non-neuronal cells

Endosomes are proposed to be the primary site of BACE-1-mediated APP cleavage (Thinakaran and Koo, 2008; Vassar et al., 1999; Morel et al., 2013; Das et al., 2013, 2015). Quantitative determination of the APP trafficking itinerary from synthesis to endosomes is challenging, which has prevented reliable measurements of APP endocytosis in neurons. To specifically visualize endocytosed APP, we designed an APP construct with an N-terminal streptavidin (SA)-binding peptide (SBP) and a C-terminal GFP (SBP-APP^wt^-GFP; **Figure 1A**). The SBP efficiently binds fluorescent SA (SA555) in the culture medium; because SA is not membrane permeant (McCann et al., 2005; Nabb and Bentley, 2022), any SA-labeled intracellular APP must have trafficked to the plasma membrane and undergone endocytosis (**Figure 1B**). This strategy was adapted from a previous study that verified the specificity of this labeling approach (Nabb and Bentley, 2022).

**Figure 1.**
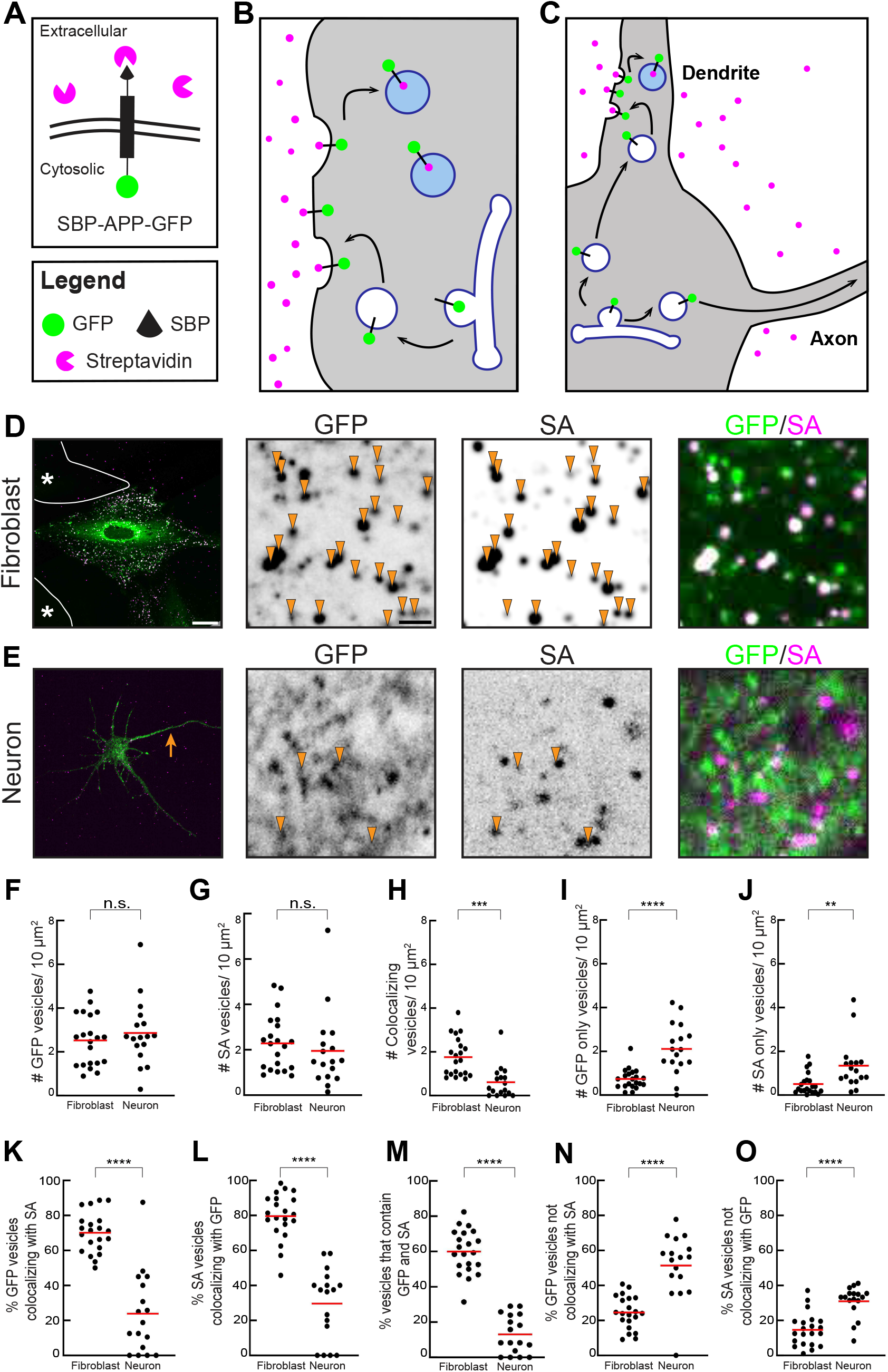
(A) A schematic illustrating the construct to differentially label Golgi-derived and endocytosed APP. APP was designed with a streptavidin-binding peptide (SBP) in the ectodomain and a cytoplasmic GFP. Cells were treated with streptavidin (SA) conjugated to Alexa Fluor 555, which cannot cross the plasma membrane. If the SBP is exposed to the medium, it binds SA with high affinity. Any vesicles labeled with SA contain APP that has trafficked via the cell surface, while vesicles labeled only with GFP have not. (B, and C) Schematics of endosomal APP labeling in an unpolarized rat embryonic fibroblast (B) or a neuron (C). (D, and E) Representative images of rat embryonic (D) or an 8 DIV hippocampal neuron (E) expressing SBP-APP^wt^-GFP and treated with SA. Asterisks indicate untransfected cells not labeled by GFP or SA. The yellow arrow indicates the axon. Arrowheads in high magnification images indicate endosomes labeled by GFP and SA. Scale bars: low mag: 20 μm, high mag: 2 μm. (F-J) Quantification of GFP and SA objects. (K-O) Quantification of GFP and SA colocalization. Red lines indicate means. Sample sizes: fibroblasts: 21 cells; neurons: 17 cells. p values: **p<0.01, ***p<0.001, ***p<0.0001.

We expressed SBP-APP^wt^-GFP in rat embryonic fibroblasts, unpolarized cells with a relatively filat profile that is conducive to high-resolution microscopy (Bentley et al., 2015; Heidemann et al., 1999; Bentley and Banker, 2015). Endocytosed APP was labeled by treating cells with SA555 for 30 min, followed by a 2 h washout, fixation, and imaging (**Figure 1D**). The Golgi exhibited the expected GFP signal from newly synthesized SBP-APP^wt^-GFP and a complete absence of SA. Vesicles were visible throughout the cell periphery. Of these, most were labeled by both GFP and SA, indicating that they contained endocytosed APP. Endosomal localization of APP was observed previously in non-neuronal cells (Koo et al., 1996; Caporaso et al., 1994), although it is unclear whether APP reaches endosomes only by endocytosis or also directly from the Golgi (Toh et al., 2017; Tam et al., 2014). We quantified the number of vesicles in the GFP and SA channels (**Figure 1F-O**). We measured 2.5 GFP, 2.3 SA, and 1.8 colocalizing vesicles per 10 µm^2^. Only 0.5 vesicles per 10 µm^2^ were labeled exclusively by GFP (**Figure 1I**) indicating that they contained APP that had not trafficked to the plasma membrane. Most APP vesicles were endosomal, with 70 % of GFP vesicles also containing SA (**Figure 1K**). These data show that most APP localizes to endosomes in unpolarized cells and that APP reaches endosomes via endocytosis and not directly from the Golgi.

Because neurons are unusual in geometry and size, and because trafficking mechanisms in neurons differ significantly from unpolarized cells, the neuronal localization of vesicle cargoes—including APP—may differ substantially from their localization in unpolarized cell types. Thus, we performed equivalent experiments with SBP-APP^wt^-GFP in primary hippocampal neurons (**Figure 1C**). Images show that GFP and SA labeling of APP vesicles in neurons differed substantially from non-neuronal cells (**Figure 1E**). Neuronal APP vesicles appeared smaller and dimmer than in non-neuronal cells. Most GFP-positive vesicles were not labeled by SA, indicating that they contained APP that had not reached the cell surface. We measured the number of GFP- and SA-containing APP vesicles (**Figure 1F-O**) and detected 2.9 GFP, 1.9 SA, and 0.6 colocalizing vesicles per 10 µm^2^. Only 24 % of GFP-positive vesicles were colabeled by SA.

Comparison of fibroblast and neuron measurements revealed significant differences in their APP distribution (**Figure 1F-O**). In non-neuronal cells, 70 % of GFP-labeled vesicles were also SA positive. This was substantially lower in neurons (24 %). Notably, most neuronal endosomes labeled by SA were not colabeled by GFP (31 %). N-terminal fragments labeled with SA remain trapped in the vesicle after APP processing; cleaved C-terminal GFP-tagged fragments are released into the cytoplasm. Therefore, SA vesicles without GFP likely indicate that APP cleavage is more efficient in neuronal endosomes. Overall, measurements consistently found significant differences in GFP and SA labeling between neurons and non-neuronal fibroblasts, highlighting the cell type-specific biology that underlies APP trafficking.

### APP^wt^ and APP^Swe^differ in their neuronal localizations

Proteolytic processing and Alzheimer’s disease progression are altered by multiple APP mutations (Hunter and Brayne, 2018). The Swedish APP variant (APP^Swe^; KM670/671NL) (Citron et al., 1992) is associated with early onset dementia in patients and increased Aβ production in animal and tissue culture models (Mullan et al., 1992; Citron et al., 1992, 1994; Shin et al., 2010; Perez et al., 1996; Johnston et al., 1994; Tambini et al., 2020). While much is known about APP^Swe^–BACE-1 interactions (Haass et al., 1995; Wang et al., 2024; Thinakaran et al., 1996), it is unclear if intracellular trafficking of APP^Swe^ and APP^wt^ differ.

To determine if the localization of APP^Swe^ differed from that of APP^wt^ in fibroblasts or neurons, we expressed SBP-APP^Swe^-GFP in each cell type (**Figure 2**).

**Figure 2.**
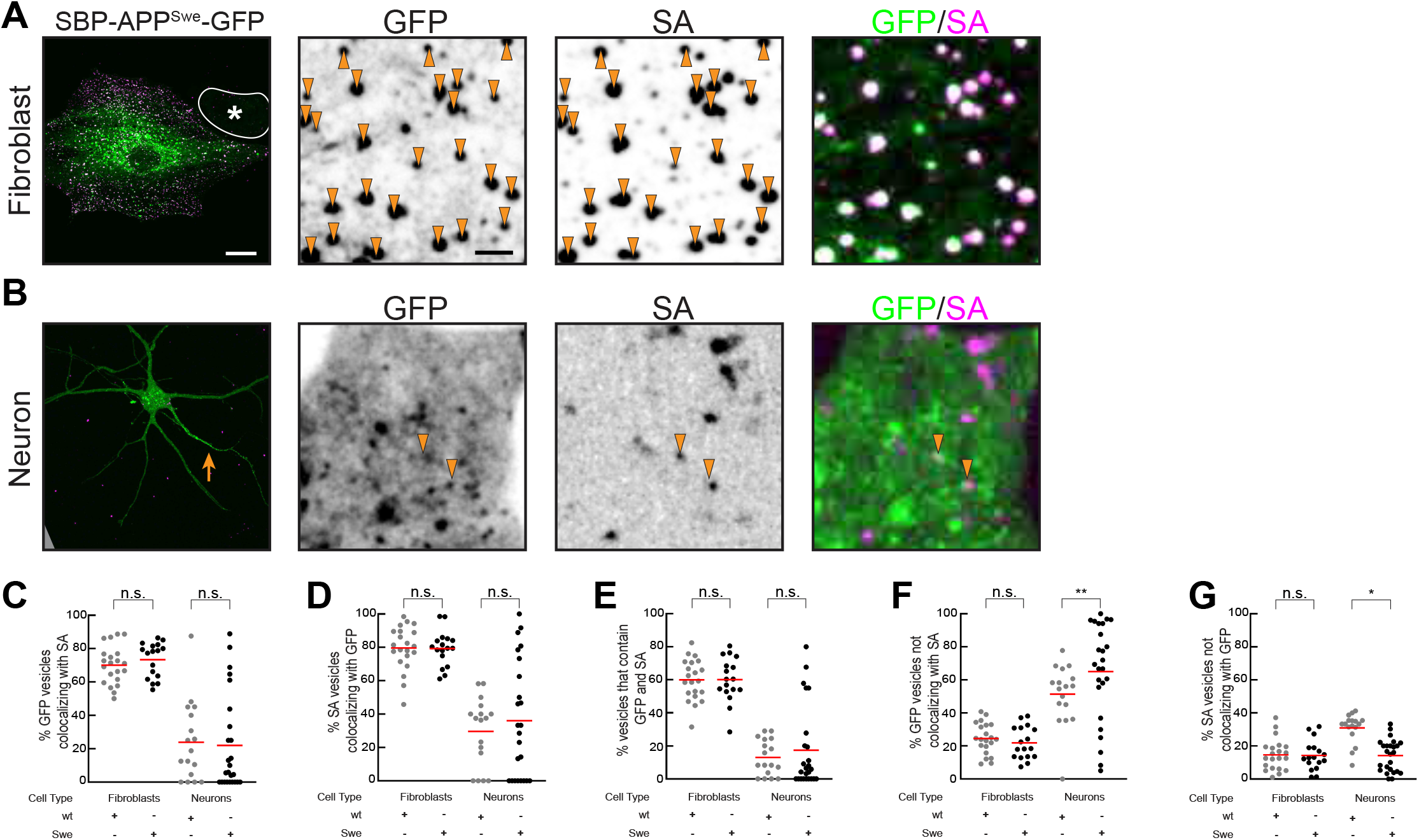
(A, and B) show representative images of a rat embryonic fibroblast (A) or a 7 DIV hippocampal neuron (B) expressing SBP-APP^Swe^-GFP and treated with SA555. Asterisk indicates an untransfected cell. The arrow indicates the axon. Arrowheads indicate endosomes labeled by GFP and SA. Scale bars: low mag: 20 μm, high mag: 2 μm. (C-G) Quantification of GFP and SA colocalization. Grey dots are data with SBP-APP^wt^-GFP and reproduced from Figure 1. Red lines show means. Sample sizes: APP^Swe^ fibroblasts: 17 cells; APP^Swe^ neurons: 24 cells. p values: *p<0.05, **p<0.01.

High magnification images show that GFP and SA signals of APP^Swe^ in fibroblasts were comparable to APP^wt^ (**Figures 1D&2A**). Most APP^Swe^ was in SA-positive endosomes. Quantifications comparing APP^wt^ and APP^Swe^ in fibroblasts showed no significant differences in their localization (**Figure 2C-F**).

In contrast to fibroblasts, there were distinct differences in the neuronal distributions of APP^wt^ and APP^Swe^ (**Figure 1E&2B**). More APP^Swe^ was in non-endosomal vesicles—likely Golgi-derived (**Figure 2F**; APP^wt^ = 51 %, APP^Swe^ = 65 %). This increase in non-endosomal APP^Swe^ compared to APP^wt^ was not due to an overall increase in endosomes, as those numbers did not differ significantly between the conditions (APP^wt^: 1.9/µm^2^ ±0.41; APP^Swe^: 1.6/µm^2^ ±0.27). These observations indicate that APP^Swe^ was not excluded from endosomes but entered additional vesicle populations that were not accessible to APP^wt^.

These data show that the residue changes in APP^Swe^ change its neuronal localization. This difference raises questions about the specific sorting locations where APP^wt^ and APP^Swe^ diverge. Because of the complexity of APP trafficking, it is possible that differences in multiple trafficking steps have synergistic effects that result in increased exposure to BACE-1 and enhanced Aβ production for APP^Swe^. In addition, the differences in APP localization between neurons and non-neuronal cells highlight the importance of defining the neuron-specific trafficking of APP^wt^ and APP^Swe^.

### APP^Swe^ enters additional post-Golgi vesicles that are not accessible to APP^wt^

The trans-Golgi network is the central hub of the secretory pathway that sorts newly synthesized transmembrane proteins. It is likely the first trafficking step where sorting of APP^wt^ and APP^Swe^ may differ. To identify Golgi-derived APP vesicles, we expressed APP^wt^-GFP or APP^Swe^-GFP with a signal sequence (SigSeq)-mCherry construct (El Meskini et al., 2001; Nabb and Bentley, 2022; Montgomery et al., 2024). This construct consists of the neuropeptide Y signal sequence fused to mCherry. It labels Golgi-derived vesicles by targeting the fluorophore into the lumen of the secretory pathway; fusion of post-Golgi vesicles with the plasma membrane results in release of the mCherry into the culture medium (Das et al., 2013; Ganguly et al., 2017; Nabb and Bentley, 2022). Any vesicles labeled by both GFP and mCherry are therefore pre-exocytic Golgi-derived vesicles that contain APP.

Low magnification images show that SigSeq-mCherry, APP^wt^-GFP, and APP^Swe^-GFP vesicles were in dendrites but were enriched in axons (**Figure 3A&B**). To evaluate the transport parameters of Golgi-derived APP^wt^ and APP^Swe^, we generated kymographs from axons and dendrites (**Figure 3C-F**). Kymograph lines with a positive slope indicate anterograde movement and those with a negative slope indicate retrograde movement, a convention followed throughout this article. Most APP^wt^ and APP^Swe^ that moved anterograde were in Golgi-derived vesicles, indicating that this was their primary pathway to the axon.

**Figure 3.**
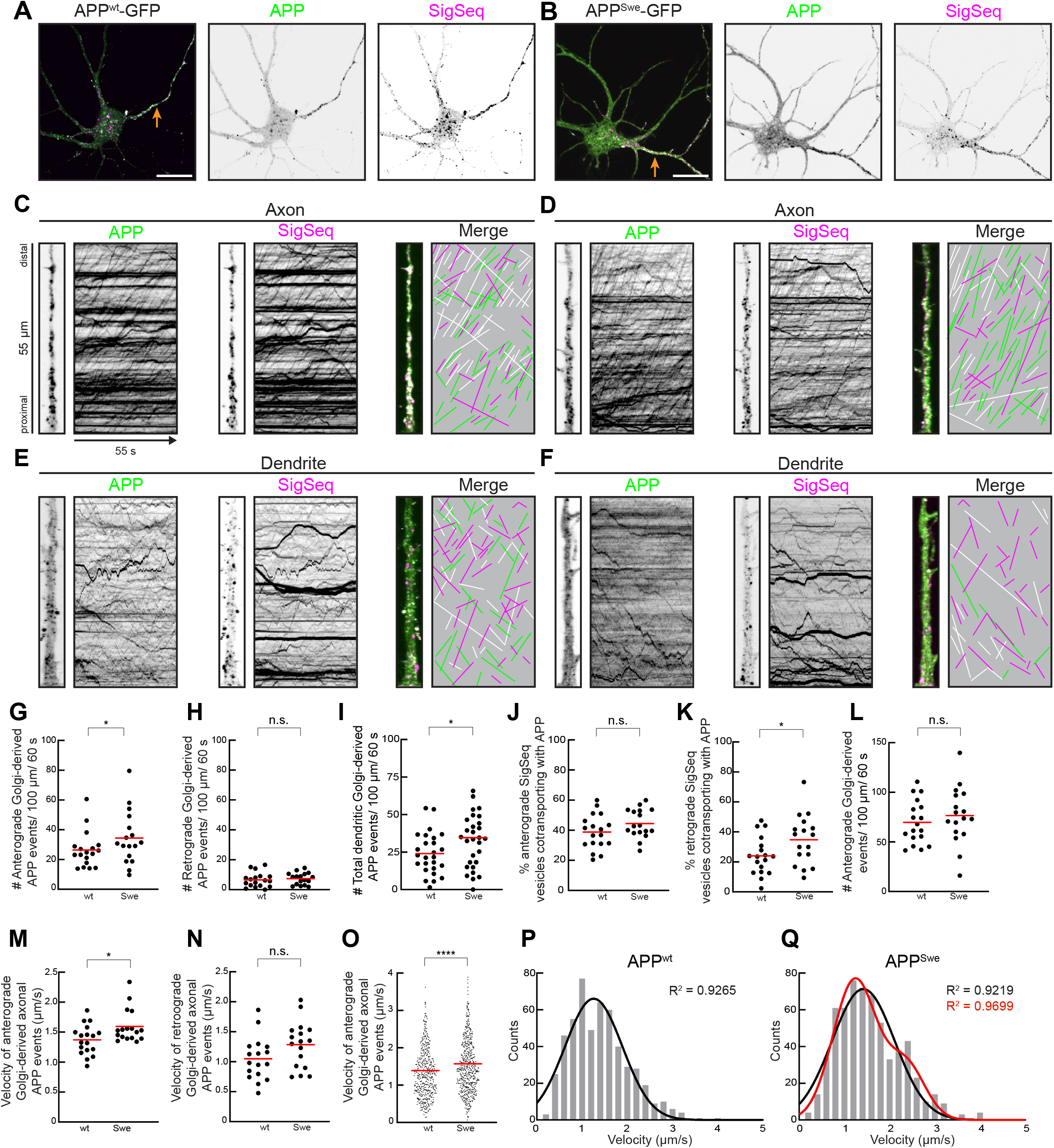
(A, and B) Representative images of 9 and 7 DIV hippocampal neurons expressing APP^wt^-GFP or APP^Swe^-GFP with SignalSequence-mCherry to label Golgi-derived vesicles. The arrows indicate axons. Scale bar: 20 μm. (C-F) Representative high magnification images and kymographs from axons and dendrites. Lines with a positive slope indicate anterograde movement, lines with a negative slope indicate retrograde movement. For clarity, transport events were redrawn. Green lines represent APP-only, red lines SigSeq-only, and white lines cotransport events. (G, H, and J-O) Quantification of cotransport in axons. Sample sizes: APP^wt^ 18 cells; APP^Swe^ 17 cells. One APP^wt^ and two APP^Swe^ data points were omitted from 3O because they exceeded the y-axis. (I) Quantification of cotransport in dendrites. Red lines show means. Sample size: APP^wt^: 26 dendrites from 11 neurons; APP^Swe^: 30 dendrites from 12 neurons. (P, and Q) Histograms showing the distribution of anterograde velocities. Black lines indicate fit to a single Gaussian distribution. The red line indicates fit to a sum of two Gaussian distributions. Sample sizes: APP^wt^: 527 events; APP^Swe^: 622 events. p values: *p<0.05, ****p<0.0001.

There were fewer APP^wt^ and APP^Swe^ transport events in dendrites than in axons, and most did not cotransport with SigSeq (**Figure 3E&F**). This supports the hypothesis that most APP endosomes are generated in the somatodendritic domain. In axons, APP^wt^-GFP and APP^Swe^-GFP exhibited primarily anterograde movements, consistent with previous reports (Szodorai et al., 2009; Fu and Holzbaur, 2013; Kaether et al., 2000; Tang et al., 2012). Both APP^wt^-GFP and APP^Swe^-GFP cotransported with SigSeq-mCherry, indicating that most axonal APP vesicles were Golgi-derived. The APP transport we observed was comparable to canonical axon-selective vesicle movement (Burack et al., 2000; Nabb and Bentley, 2022; Frank et al., 2022; Petersen et al., 2014; Nabb et al., 2020) and consistent with APP being primarily targeted to axons (Ferreira et al., 1993; Yamazaki et al., 1995). It is possible that APP transports in some of the same vesicles that move axonally polarized membrane proteins.

Because most newly synthesized APP is targeted to the axon, we quantified axonal transport of Golgi-derived APP^wt^ or APP^Swe^ vesicles. APP^Swe^-GFP trafficked in more Golgi-derived vesicles undergoing anterograde movement than APP^wt^-GFP (**Figure 3G**). In contrast, there was no difference in the retrograde movements (**Figure 3H**). Similarly, there were more Golgi-derived APP^Swe^ vesicles in dendrites than APP^wt^ (**Figure 3I**).

The larger number of APP^Swe^ post-Golgi vesicles could be for two reasons: either an overall increase in the formation of Golgi-derived vesicles, or because APP^Swe^ was sorted less specifically at the trans-Golgi and entered additional constitutive post-Golgi vesicles. To differentiate these models, we measured the percentage of Golgi-derived (SigSeq-mCherry) vesicles that cotransported with APP (**Figure 3J&K**). APP^Swe^ was found in a larger fraction of SigSeq vesicles, although this difference was only statistically significant in retrograde movements. Because the total number of SigSeq vesicles was unaffected (**Figure 3L**), this indicates that APP^Swe^ expression did not result in additional post-Golgi vesicles, but that APP^Swe^ entered a larger fraction of the available post-Golgi vesicle pool. Therefore, APP^Swe^ sorting at the trans-Golgi is less specific than sorting of APP^wt^. That could be because APP^Swe^ has lost a sorting signal and is sorted less stringently than APP^wt^. Alternatively, APP^Swe^ may have acquired a new sorting signal that actively targets some of it into additional and distinct post-Golgi vesicles.

To determine if APP^Swe^ was sorted into an additional population of Golgi-derived vesicles with distinct transport parameters, we analyzed the velocities of Golgi-derived APP^wt^ and APP^Swe^ vesicles (**Figure 3M-O**). Anterograde vesicle transport is mediated by some 20 kinesins (Silverman et al., 2010). Because each vesicle population is moved by a specific complement of kinesins, there are transport *signatures* (e.g., velocity), that correlate with a particular vesicle identity (Yang et al., 2019; Garbouchian et al., 2022; Frank et al., 2020, 2022). Anterograde velocities of APP^wt^ were lower than for APP^Swe^ (**Figure 3M**). Retrograde velocities did not differ significantly, presumably because all long-range retrograde movement in axons is mediated by dynein (**Figure 3N**). The mean velocity of all observed APP^wt^ Golgi-derived transport events was significantly lower than for APP^Swe^ (**Figure 3O**). We analyzed the distribution of anterograde transport velocities by generating histograms for Golgi-derived APP^wt^ and APP^Swe^ events (**Figure 3P&Q**). APP^wt^ velocities exhibited a typical Gaussian distribution with a single peak at 1.26 µm/s. APP^Swe^ velocities exhibited a more complex distribution. There was a primary peak at 1.23 µm/s and a smaller second peak at 2.37 µm/s. We tested fitting the APP^Swe^ velocity histogram data to a Gaussian (R^2^ = 0.9219) or sum of two Gaussian (R^2^ = 0.9699) distributions. We compared fits with an Akaike Information Criterion to determine the quality of the two models and found that sum of two Gaussian distribution outperformed the single Gaussian.

The differences in Golgi-derived trafficking between APP^wt^ and APP^Swe^ are consistent with a model in which APP^Swe^ is sorted into at least one additional vesicle population at the trans-Golgi, indicating a possible sorting defect. Aβ generation requires APP to interact with active BACE-1, which only occurs in a subset of organelles (Das et al., 2015). These observations suggest the hypothesis that enhanced Aβ generation by APP^Swe^ may be caused by sorting of APP^Swe^ into additional Golgi-derived vesicles that contain active BACE-1.

### Dendrite to axon transcytosis is higher for APP^Swe^ than APP^wt^

Most APP reaches the axon by direct delivery from the trans-Golgi (**Figure 3C&D**), similar to other axonal membrane proteins (Nabb and Bentley, 2022). Some axonal proteins can reach the axon by transcytosis (Nabb and Bentley, 2022). In the transcytotic pathway, proteins leave the trans-Golgi in vesicles destined to the dendritic plasma membrane, where they are endocytosed and sorted into axon-selective vesicles that deliver them to the axon. Because BACE-1 enzymatic activity requires a low pH (Vassar et al., 1999), which is found in endosomes, transcytotic vesicles have been proposed as a key site for Aβ generation (Woodruff et al., 2016; Koo et al., 1996; Norstrom et al., 2010; Das et al., 2015, 2013; Niederst et al., 2015).

To determine if the rate of transcytosis between APP^wt^ and APP^Swe^ differed, we performed a live imaging time course of SBP-APP^wt^-GFP or SBP-APP^Swe^-GFP after a 30 min SA647 treatment (**Figure 4**). Axon kymographs from the SA647 channel show that there were few transcytotic events (i.e., anterograde moving vesicles) within 1 h of treatment for both APP^wt^ and APP^Swe^ (**Figure 4A&B**). This is consistent with observations with other axonal membrane proteins as it takes time for cargoes to move through the endocytic pathway (Nabb and Bentley, 2022). At 6 h after treatment, there was transcytosis of both APP^wt^ and APP^Swe^ (**Figure 4C&D**). Plots of the average number of transcytotic events show transcytosis appeared at 150 min after SA647 treatment in both conditions (**Figure 4E-G**). Transcytosis peaked 390 min after SA647 treatment. It was more pronounced in neurons expressing APP^Swe^, where the median was nearly double that of APP^wt^ (APP^wt^ = 3.38; APP^Swe^ = 6.61).

**Figure 4.**
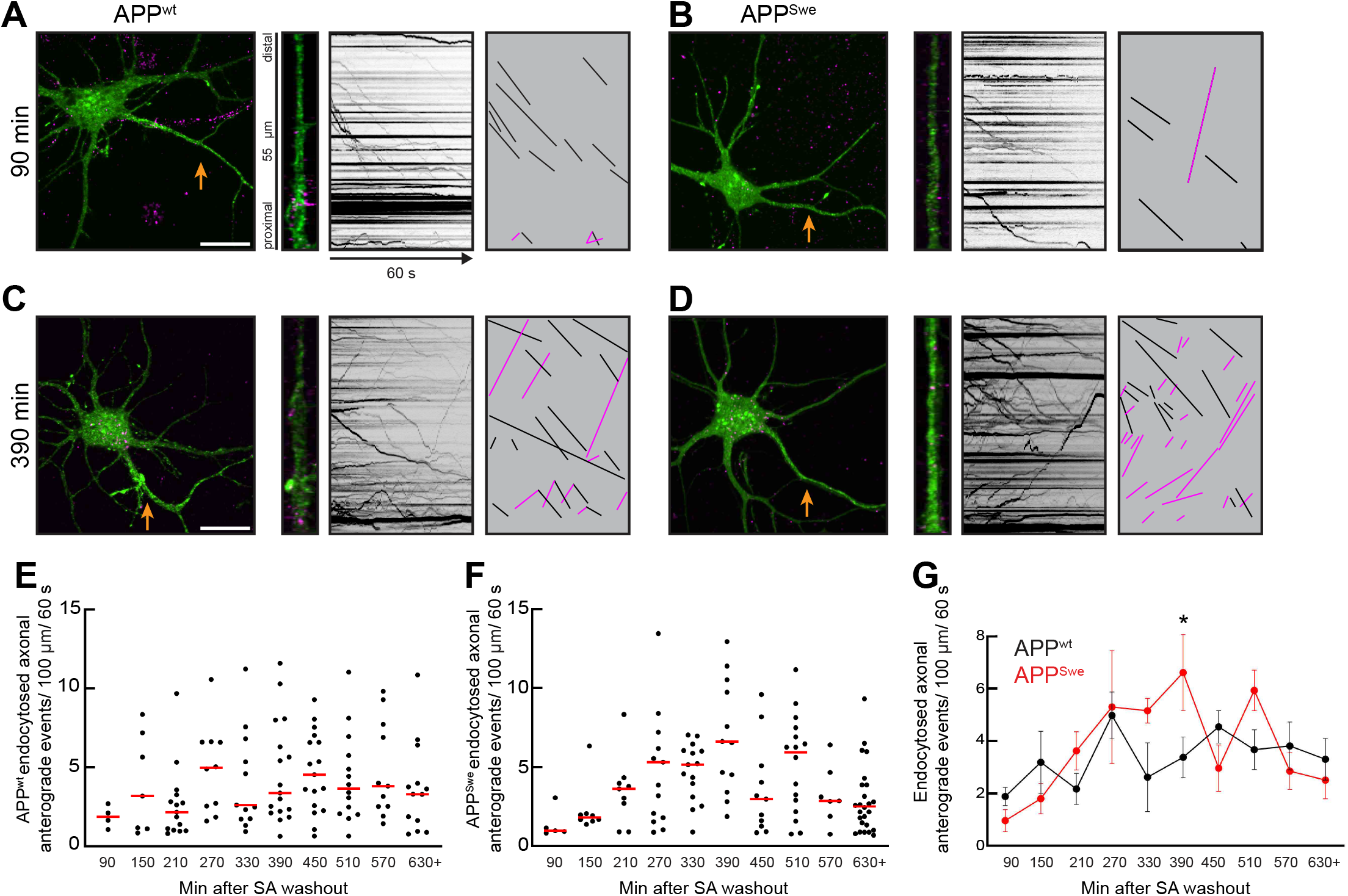
(A-D) Representative images and axon kymographs of 7 DIV neurons imaged at 1 h (A, and B) or 6 h (C, and D) after SA647 washout. Arrows indicate axons. Black lines indicate retrograde transport; magenta lines indicate anterograde transport. Scale bar: 20 μm. (E-G) Quantification of SA transport in axons. Cells measurements were assigned to 60 min bins based on the time of imaging after SA treatment. One APP^wt^ data point was omitted at 330 min for exceeding the y-axis. Four APP^Swe^ data points were omitted for exceeding the y-axis; two at 270 min, one at 360 min, and one at 690+ min. Red lines show medians. Sample sizes for each bin (APP^wt^/APP^Swe^): 90 (4/5), 150(7/8), 210 (15/9), 270 (10/15), 330 (13/15), 390 (17/13), 450 (17/11), 510 (14/16), 570 (11/7), 630 (13/26). p value: *=0.04.

These experiments show that APP^Swe^ is more likely to reach the axon in transcytotic vesicles than APP^wt^. However, it is unclear if this difference in transcytosis is attributable to only the previously found differences in post-Golgi trafficking. The additional APP^Swe^ Golgi-derived vesicles may be more likely to traffic APP^Swe^ to the dendritic plasma membrane, resulting in more dendritically endocytosed APP^Swe^ and increased transcytosis. Alternatively, greater APP^Swe^ transcytosis could also be caused by differences in sorting between APP^wt^ and APP^Swe^ after endocytosis, with APP^Swe^ being more likely to enter transcytotic vesicles.

### Endocytosed APP^Swe^ is targeted to lysosomes less efOiciently than APP^wt^

Endocytosed APP that is not recycled to the dendritic membrane can be targeted to lysosomes (Haass et al., 1992; Lai et al., 1995; Caporaso et al., 1992; Lorenzen et al., 2010) or travel to the axon by transcytosis (Woodruff et al., 2016). Experiments above found that, compared to APP^wt^, a smaller fraction of endocytosed APP^Swe^ vesicles in the cell body were not labeled by GFP (**Figure 2G**). Loss of GFP is associated with successful γ-secretase cleavage. This observation is consistent with endocytosed APP^Swe^ undergoing enhanced transcytosis to the axon. Increased transcytosis could be due to less efficient sorting of endocytosed APP^Swe^ to lysosomes.

To determine if endocytosed APP^Swe^ is more likely to avoid lysosomal targeting, we expressed SBP-APP^wt^-GFP or SBP-APP^Swe^-GFP, treated cells with SA647, and applied lysotracker to label acidified lysosomes (**Figure 5**). Neurons were recorded at 1 h or 2 h post SA washout to track APP through the endocytic pathway (**Figure 5A-D**). We chose these time points to account for the time it takes for dendritically endocytosed APP to traffic to the soma and reach lysosomes. At both time points after washout, we observed the expected SA labeling in somata. Quantification found that the number of endocytosed APP^wt^ vesicles that had not reached lysosomes was larger than for APP^Swe^ vesicles, although not statistically significant (p values: 60 min=0.051; 120 min=0.055) (**Figure 5E&F**). The rate of APP delivery to lysosomes, as determined by the number of lysosomes that had not acquired endocytosed APP, was consistent for both APP^wt^ and APP^Swe^ (**Figure 5G&H**). Consolidation of endocytosed APP into lysosomes is expected to result in a decrease of total SA-positive vesicles between 60 and 120 min, which we observed for both conditions (**Figure 5I&J**). A second potential pathway to decrease of pre-lysosomal endosomes could be transcytosis of endosomes to axons. We observed a greater decrease in the number of endocytosed vesicles for APP^Swe^ than APP^wt^. Because the overall number of lysosomes was consistent between conditions, the exacerbated loss of vesicles in the 120 min APP^Swe^ condition suggests an additional pathway by which these vesicles were depleted, potentially by increased transcytosis. These observations were consistent with endosomes containing APP^wt^ primarily reaching the soma and being targeted to lysosomes, whereas a greater subset of endocytosed APP^Swe^ vesicles traversed the soma and entered the axon, consistent with the finding that APP^Swe^ was more likely to undergo transcytosis (**Figure 4G**). These data also indicate that sorting of APP^wt^ and APP^Swe^ differs not just at the trans-Golgi, but also after dendritic endocytosis.

**Figure 5.**
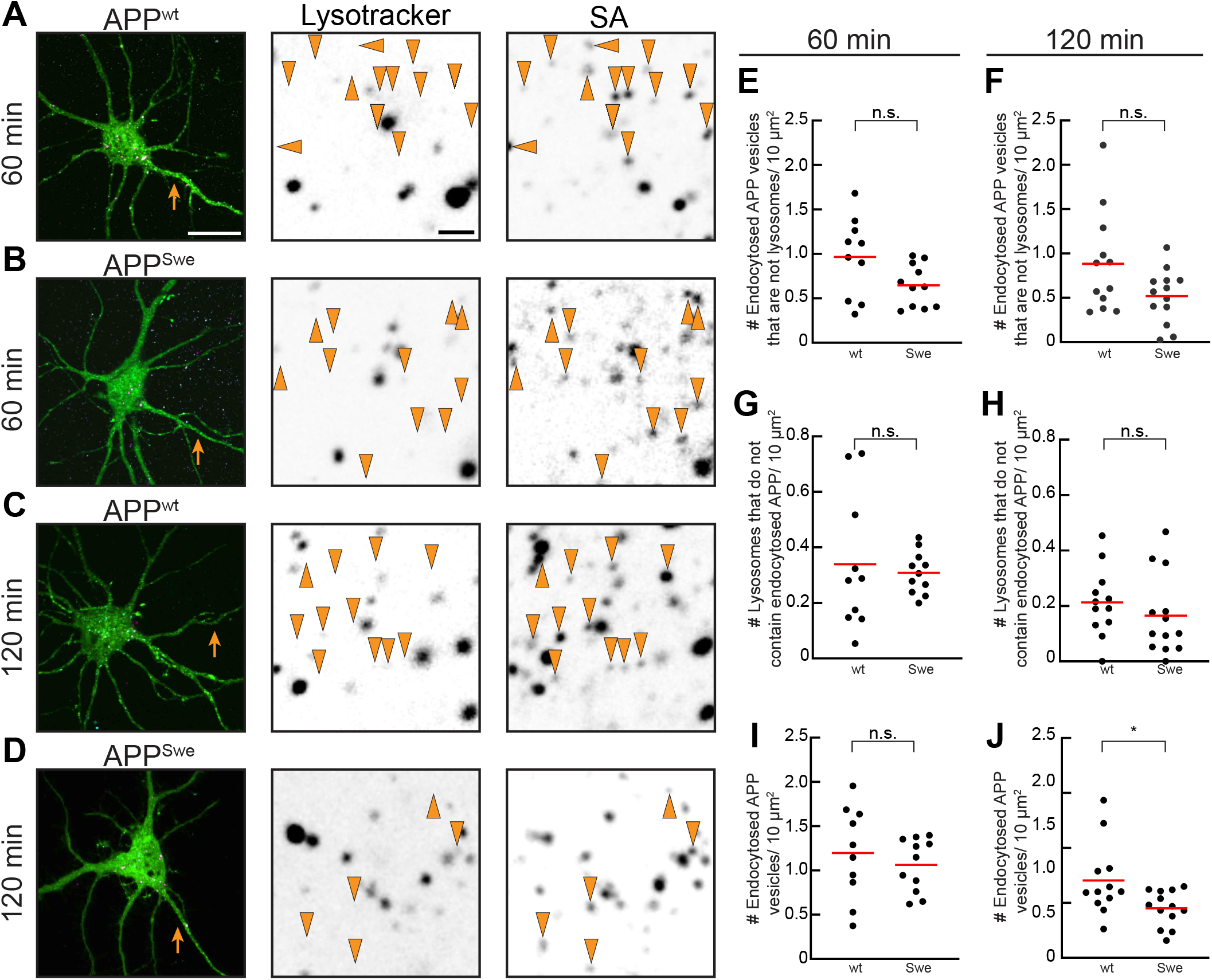
(A-D) Representative images of 7-8 DIV neurons expressing SBP-APP^wt^-GFP or SBP-APP^Swe^-GFP and treated with 19.5 nM SA647 and 10 nM lysotracker red. Arrows indicate axons. Arrowheads on high magnification somata images indicate endosomes that did not colocalize with lysotracker. Red lines show means. Scale bars: low mag: 20 μm, high mag: 2 μm. (E-J) Quantification of SA and lysotracker objects. Sample sizes (APP^wt^/APP^Swe^): 90 min (10/11), 120 min (12/13)).

### APP^Swe^increases Aβ formation in Golgi-derived and transcytotic vesicles

BACE-1-mediated APP cleavage and subsequent Aβ formation are prominent components of AD progression (Thinakaran and Koo, 2008; O’Brien and Wong, 2011). Experiments with non-neuronal cells found that BACE-1 cleaves APP in the secretory pathway (Tan and Gleeson, 2019b; Greenfield et al., 1999; Thinakaran et al., 1996; Xia et al., 2000; Xu et al., 1997). Data from neurons show that Aβ formation occurs in endosomes (Das et al., 2015, 2013; Woodruff et al., 2016). APP and BACE-1 cotransport in Golgi-derived vesicles (Das et al., 2015), but it is unclear if APP in the secretory pathway of primary neurons undergoes efficient BACE-1 cleavage.

To visualize BACE-1-cleaved APP, we used the VU-17 antibody, which specifically recognizes the first six residues of the cleaved Aβ N-terminus (Verwey et al., 2013). To verify the specificity of the antibody, we stained neurons expressing human APP-GFP with VU-17 and quantified the staining (**Figure 6A&B**). VU-17 is specific for human APP, and untransfected cells exhibited non-specific background staining. Transfected neurons expressing APP-GFP had punctate VU-17 staining. Treatment with the BACE-1 inhibitor verubecestat (Kennedy et al., 2016) resulted in a sharp decrease in staining, demonstrating that VU-17 labeling was contingent on BACE-1 activity and an indicator of APP that has been targeted for amyloidogenesis.

**Figure 6.**
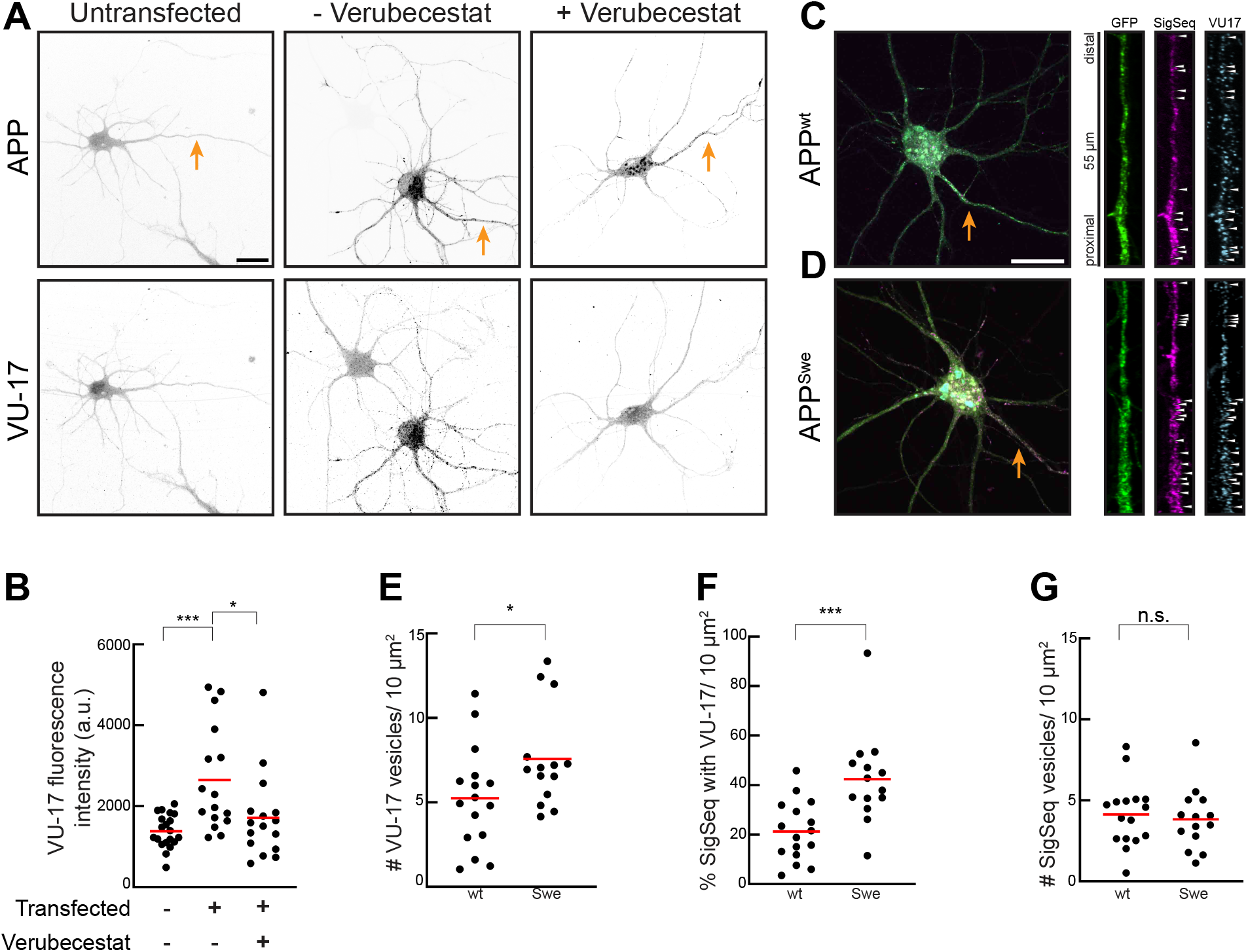
(A) Representative images of 7 DIV untransfected neurons or neurons expressing APP^wt^-GFP in the presence and absence of the BACE-1 inhibitor verubecestat and stained with VU-17 which recognizes the N-terminus of cleaved Aβ. (B) Quantification of VU-17 staining. Sample sizes (transfected/verubecestat): (-/-) 20; (+/-) 16; (+/+) 16. (C, and D). Representative images 7 or 8 DIV neurons expressing APP^wt^-GFP or APP^Swe^-GFP with SignalSequence-mCherry and stained with VU-17. Arrows indicate axons. High magnification images show the axon of each neuron. Arrowheads indicate colocalization between SigSeq and VU17. Scale bars: 20 μm. (E) Quantification of VU-17 objects. Red lines show means. (F) Quantification of SigSeq and VU-17 colocalization. (G) Quantification of SigSeq objects. Sample sizes: APP^wt^: 16 cells; APP^Swe^: 14 cells. p values: *p<0.05, ***p<0.001..

To determine if there were differences in the amyloidogenic processing of APP^wt^ and APP^Swe^ in Golgi-derived vesicles, we co-expressed APP^wt^-GFP or APP^Swe^-GFP with SigSeq-mCherry and stained with VU-17 (**Figure 6C&D**). High magnification images of axons show specific VU-17 labeling of neurons expressing APP^wt^ and APP^Swe^. Most of the staining colocalized with SigSeq, indicating that these BACE-1 cleavage products were localized to Golgi-derived vesicles. fuantification found that there were more VU-17 structures in axons from neurons expressing APP^Swe^ than APP^wt^ (**Figure 6E**). We determined the fraction of axonal SiqSeq vesicles that stained for VU-17 and found that a significantly higher subset of APP^Swe^ SigSeq vesicles were VU-17 positive than APP^wt^ (**Figure 6F**). This difference was not due to the formation of more SigSeq vesicles, as the absolute number of SigSeq vesicles was consistent between the two conditions (**Figure 6G**). These results are consistent with a model in which APP^Swe^ enters a larger subset of Golgi-derived vesicles than APP^wt^, and that these additional compartments are conducive to BACE-1 cleavage (**Figure 3**). These results show that BACE-1-mediated cleavage of APP is not limited to endosomes. Instead, APP^wt^ and APP^Swe^ are both subject to cleavage in the neuronal secretory pathway.

## DISCUSSION

This study made three principal discoveries. First, targeting of APP to endosomes is more efficient in unpolarized cells than in primary neurons. Second, APP^wt^ and APP^Swe^ have the same localization in unpolarized cells but exhibit different trafficking in neurons. APP^Swe^ is sorted into additional vesicles at the trans-Golgi and more likely to be sorted into transcytotic vesicles after dendritic endocytosis. Third, APP cleavage by BACE-1 is not restricted to neuronal endosomes but also common in Golgi-derived vesicles.

These findings show that differences in the trafficking of APP^wt^ and APP^Swe^ may be key contributors of enhanced Aβ production associated with APP^Swe^. They also show that there are differences in APP trafficking between neurons and unpolarized cells and that understanding neuronal APP trafficking and Aβ production necessitates experiments in neurons.

### TrafOicking differences between APP^wt^ and APP^Swe^

We identified two trafficking events at which APP^wt^ and APP^Swe^ are processed differently: at the trans-Golgi and during endocytosis at the dendritic plasma membrane. Both sorting events have been linked to heterotetrameric clathrin adaptor complexes (AP) (Bonifacino, 2014). Golgi exit of APP is mediated by AP-1 and AP-4 (Burgos et al., 2010; Januário et al., 2022; Icking et al., 2007) and endocytosis by AP-2 through interaction with X11 (Borg et al., 1996) or Dab2 (Lee et al., 2008; Nordstedt et al., 1993). In each case, sorting has been mapped to YXXØ motifs in the APP cytoplasmic domain. This is consistent with our data, as most APP^Swe^ entered the same vesicles as APP^wt^, and a small fraction of APP^Swe^ appeared in non-APP^wt^ vesicles. It is notable that the sorting difference is caused by residue changes in the APP ectodomain. Another neuronal type I membrane protein, NgCAM/L1CAM, relies on signals in its ectodomain to reach its proper distribution (Sampo et al., 2003), despite also interacting with AP complexes through a YXXØ motif (Yap et al., 2008; Kamiguchi et al., 1998).

The data here show that multiple signals—in the ecto- and cytoplasmic domains of APP—contribute to APP sorting. The KM670/671NL change in APP^Swe^ may cause sorting changes in at least two ways. First, the NL mutation could act as a second sorting motif that actively sorts APP into different vesicles; second, loss of the KM residues could be the loss of a specific sorting signal and degrade the fidelity of the sorting. The data in this study cannot differentiate between these models. In general, sorting motifs in the luminal domains of cargo proteins are much less defined than cytoplasmic motifs (Watson et al., 2025), particularly in neurons (Bentley and Banker, 2016).

In addition to the two trafficking stages where APP^wt^ and APP^Swe^ are sorted differently, there are further potential events of trafficking divergence. There is evidence of retromer-mediated trafficking of APP (Nielsen et al., 2007; Schmidt et al., 2007; Andersen et al., 2005), and APP may undergo retrograde trafficking from endosomes to the Golgi. A Golgi-to-endosome pathway has also been proposed (Toh et al., 2017). If APP^wt^ and APP^Swe^ diverge at these trafficking events, remains to be seen.

### APP trafOicking and Aβ formation

If and how APP trafficking determines Aβ generation is a long-standing and unresolved problem (Sun and Roy, 2018; Tan and Gleeson, 2019a). A strength of this study is the systematic comparison of specific compartments within one relevant model system, primary hippocampal neurons. That approach helps contextualize previous studies that focused on specific compartments or were performed in other model systems.

Combining tools to specifically visualize defined APP vesicle populations (i.e., Golgi-derived and endosomal vesicles) with immunostaining for BACE-1-cleaved APP, revealed that Aβ generation occurred in the secretory pathway and in endosomes. Our results build on previous findings that APP and BACE-1 cotransported in both compartments (Das et al., 2015) and show that APP–BACE-1 interactions in the secretory pathway result in productive cleavage and Aβ formation. APP processing in endosomes is a target for therapy development (Sun et al., 2019). Our data suggest that the neuronal secretory pathway may be an additional promising target for attenuating Aβ formation in neurons. There is evidence that APP from different parts of the endomembrane system may converge in several compartments: Golgi-derived APP may traffic directly to endosomes; endocytosed APP may travel retrograde to the trans-Golgi. The prevalence of these trafficking steps in neurons, and how such compartmental crosstalk affects Aβ formation, remain unclear. Finally, APP trafficking can change dynamically during development (Ramaker et al., 2013, 2016), which has not been explored sufficiently in mammalian neurons (Copenhaver and Ramaker, 2016).

### Neuronal transport mechanisms of APP

Most anterograde vesicle transport in neurons is mediated by kinesins, which move towards the plus ends of microtubules (Cason and Holzbaur, 2022; Yildiz, 2024). Axonal transport of APP is thought to be mediated by Kinesin-1 family members (Ferreira et al., 1993; Kamal et al., 2000) through interactions between the kinesin light chain and c-Jun N-terminal kinase-interaction protein 1 (Fu and Holzbaur, 2013; Taylor et al., 2016; Matsuda et al., 2003; Chiba et al., 2014, 2017). In aggregate, this is strong evidence that Kinesin-1 family members mediate some transport of APP vesicles.

It is possible that Kinesin-1s are not the exclusive APP transporters. Neurons express some 20 kinesins that mediate vesicle transport and may participate in the movement of various APP subpopulations (Silverman et al., 2010; Yang et al., 2019; Nabb et al., 2020) as APP is transported in many biochemically distinct vesicle types. In fact, data in this study show that APP^Swe^ moves in at least two different Golgi-derived vesicle populations with different transport parameters. Because the velocities of these vesicles differ, it is likely that their kinesins also differ. As understanding of kinesin–vesicle interactions has grown, and as more tools have become available to determine kinesin biology in a cellular context (Montgomery et al., 2022; Yang et al., 2016; Garbouchian et al., 2022; Bentley and Banker, 2015; Jenkins et al., 2012), the time may be ripe to reevaluate the kinesins that move each APP vesicle type and determine the network of kinesins that ensures the transport and processing of APP in its various cellular compartments.

## ACKNOWLEDGMENTS

We thank members of our lab group for their helpful comments on early drafts of the manuscript. We acknowledge the BioResearch facility at Rensselaer for assistance with husbandry and tissue collection. This work was supported by National Institutes of Health grant R35GM153259 to M.B. and an NIA Training Program T32AG078123 pre-doctoral fellowship to C.R.P.

## MATERIALS AND METHODS

### Cell culture

Rat embryonic fibroblasts (Heidemann et al., 1999; Bentley et al., 2015) were grown at 37°C in Dulbecco’s modified Eagle’s medium (Corning) containing 10 % fetal bovine serum, 4.5 g/l d-glucose, 548 mg/l L-glutamine, and 110 mg/l sodium pyruvate. Cells were trypsinized, plated on glass coverslips, and incubated at 37°C with 5 % CO^2^ overnight prior to transfection. Fibroblasts were transfected with Lipofectamine 2000 (Thermo Fisher Scientific, Catalog# 11668019). Cells were fixed with 4 % paraformaldehyde /4 % sucrose in phosphate-buffered saline (PBS) for 20 min at 37°C after 24 h expression.

Primary hippocampal neurons were cultured as previously described (Kaech and Banker, 2006; Kaech et al., 2012a). E18 rat hippocampi were dissected, trypsinized, dissociated, and plated onto 18 mm glass coverslips coated with poly-L-lysine. Cultured neurons were grown in N2-supplemented MEM and maintained at 37°C with 5 % CO^2^. Stage 4/5 hippocampal neurons were transfected with Lipofectamine 2000 (Thermo Fisher Scientific, cat# 11668019). The following expression times were used: Figures 1-3: 17-20 h, Figure 4: 17–28 h, Figure 5: 17-23, Figure 6: 18 h. Each experiment contains data from multiple separate transfections and at least two independent cultures.

### DNA constructs

Construct details are described in Table 1. All constructs were verified by sequencing.

### ImmunoOluorescence

Neurons were fixed with 4 % paraformaldehyde/0.4 % sucrose in PBS for 20 min at 37°C and permeabilized with 0.5 % Triton X-100 in PBS. Neurons were incubated with 0.5 % fish skin gelatin in PBS for at minimum 1 h to block nonspecific antibody-binding sites. Coverslips were incubated with primary antibody for 1 h, washed with PBS, incubated with secondary antibody for 30 min, and mounted on slides with Prolong Diamond Anti-fade (Thermo Fisher Scientific cat# P36965). BACE-1-cleaved human APP was detected with Amyloid-beta (N-term), Human, mAb VU-17 (Hycult Biotech, Cat: HM2325). To determine antibody efficacy, VU-17 signal was quantified by analyzing the brightest 10% of pixels within the cell.

### SA and lysotracker labeling

Neurons were incubated for 30 min in 12-well plates with conditioned medium and 19.5 nM of SA conjugated to Alexa Fluor 555 (Thermo Fisher, cat# S21381) or Alexa Fluor 647 (Thermo Fisher, cat# S21374) as indicated in the figure legends. After a 30 min incubation, unbound SA was removed by returning cells to SA-free medium until the cells were imaged. For lysotracker experiments, 10 nM lysotracker deep red (Thermo Fisher, cat# L12492) was added to incubating cells 5 min before rinsing. Washed coverslips were then loaded into a prewarmed imaging chamber, submerged in Hibernate E medium without phenol red (Brain-Bits), and transferred to the microscope for imaging.

### Imaging

All images and movies were acquired with an Andor Dragonfly built on a Ti2 (Nikon) with a CFI Apo 60x 1.49 objective (Nikon) and two sCMOS cameras (Zyla 4.2+; Andor). The imaging stage, microscope objectives, and cell sample (live or fixed) were maintained at 37°C in a warmed enclosure (full lexan incubation ensemble; OkoLab). The z-axis movement was controlled by the Perfect Focus System on the Ti2 microscope (Nikon). Axons were identified by morphology. All live-cell recordings (Figures 3, 4, and 5) were acquired at two frames per second. Cells were maintained in Hibernate E without phenol red (BrainBits; cat# HELF500).

### Analysis

Analysis was performed by a single blinded reviewer (C.P.), except Figure 3 (S.T.). To ensure consistency and reduce bias, multiple experiments were combined into a large data set, and the analyst was blinded to the condition.

MetaMorph Image Analysis Software (Molecular Devices) was used to generate kymographs of vesicle transport. Transport events were identified on kymographs where each continuous line with a consistent slope was scored as a single transport event. A single vesicle could undergo multiple transport events if there was a distinct pause between them. Event coordinates were exported to Microsoft Excel for analysis. To ensure only microtubule-based long-range transport events were included in the analysis, excursions < 3 μm were excluded. Transport analysis in Figure 4 selected for long-range transport and only included events >10 µm.

Cotransport (Figure 3) was determined by kymograph analysis. Each channel’s transport events were first determined separately by drawing event lines on kymographs. Then, cotransport events were identified by overlaying lines from both channels. The proximal 5 µm of the axon were excluded from analysis to exclude the axon initial segment. The number of events per 100 µm was defined as the total number of events recorded, normalized to the total length of neurite analyzed for all cells in a treatment. The cotransport percentage was calculated as the number of overlapping events divided by the total number of events.

For quantification of transcytosis, the first 25 µm of the axon were excluded to select for long range transcytotic vesicle transport that had passed through the axon initial segment.

SA and lysotracker object counting analyses were performed blinded in neuron somata or cell bodies of fibroblasts and the Golgi and nuclei were excluded (Figures 1,2, and 5). VU-17 staining object counting analyses were performed in axons (Figure 6).

All statistical analyses were performed in Microsoft Excel or GraphPad Prism. All plots were generated in GraphPad Prism. The p values reported in Figures were generated by Student’s t test, equal variances.

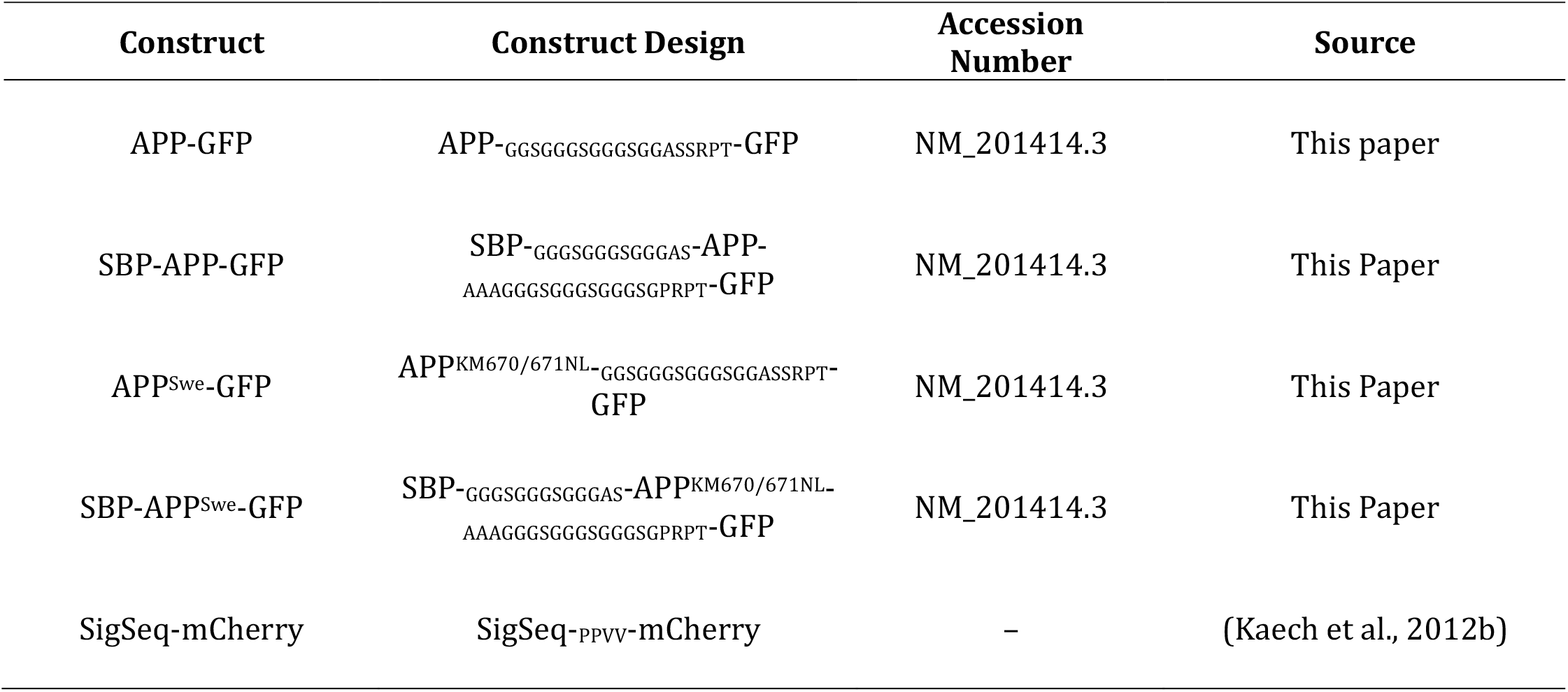

## REFERENCES

Andersen, O.M., J. Reiche, V. Schmidt, M. Gotthardt, R. Spoelgen, J. Behlke, C.A.F. Von Arnim, T. Breiderhoff, P. Jansen, X. Wu, K.R. Bales, R. Cappai, C.L. Masters, J. Gliemann, E.J. Mufson, B.T. Hyman, S.M. Paul, A. Nykjsær, and T.E. Willnow. 2005. Neuronal sorting protein-related receptor sorLA/LR11 regulates processing of the amyloid precursor protein. Proc. Natl. Acad. Sci. U. S. A. 102:13461–13466. doi:10.1073/pnas.0503689102.

Bentley, M., and G. Banker. 2015. A Novel Assay to Identify the Trafficking Proteins that Bind to Specific Vesicle Populations. Curr. Protoc. Cell Biol. 69:13.8.1-13.8.12. doi:10.1002/0471143030.cb1308s69.

Bentley, M., and G. Banker. 2016. The cellular mechanisms that maintain neuronal polarity. Nat. Rev. Neurosci. 17:611–622. doi:10.1038/nrn.2016.100.

Bentley, M., H. Decker, J. Luisi, and G. Banker. 2015. A novel assay reveals preferential binding between Rabs, kinesins, and specific endosomal subpopulations. J. Cell Biol. 93:4604. doi:10.1083/jcb.201408056.

Bonifacino, J.S. 2014. Adaptor proteins involved in polarized sorting. J. Cell Biol. 204:7–17. doi:10.1083/jcb.201310021.

Borg, J.P., J. Ooi, E. Levy, and B. Margolis. 1996. The phosphotyrosine interaction domains of X11 and FE65 bind to distinct sites on the YENPTY motif of amyloid precursor protein. Mol. Cell. Biol. 16:6229–6241. doi:10.1128/MCB.16.11.6229.

Burack, M.A., M.A. Silverman, and G. Banker. 2000. The role of selective transport in neuronal protein sorting. Neuron. 26:465–472. doi:10.1016/S0896-6273(00)81178-2.

Burgos, P.V., G.A. Mardones, A.L. Rojas, L.L.P. daSilva, Y. Prabhu, J.H. Hurley, and J.S. Bonifacino. 2010. Sorting of the Alzheimer’s Disease Amyloid Precursor Protein Mediated by the AP-4 Complex. Dev. Cell. 18:425–436. doi:10.1016/j.devcel.2010.01.015.

Caporaso, G.L., S.E. Gandy, J.D. Buxbaum, and P. Greengard. 1992. Chloroquine inhibits intracellular degradation but not secretion of Alzheimer beta/A4 amyloid precursor protein. Proc. Natl. Acad. Sci. U. S. A. 89:2252–2256. doi:10.1073/pnas.89.6.2252.

Caporaso, G.L., K. Takei, S.E. Gandy, M. Matteoli, O. Mundigl, P. Greengard, and P. De Camilli. 1994. Morphologic and biochemical analysis of the intracellular trafficking of the Alzheimer beta/A4 amyloid precursor protein. J. Neurosci. Off. J. Soc. Neurosci. 14:3122–3138. doi:10.1523/JNEUROSCI.14-05-03122.1994.

Cason, S.E., and E.L.F. Holzbaur. 2022. Selective motor activation in organelle transport along axons. Nat. Rev. Mol. Cell Biol. 23:699–714. doi:10.1038/s41580-022-00491-w.

Chiba, K., M. Araseki, K. Nozawa, K. Furukori, Y. Araki, T. Matsushima, T. Nakaya, S. Hata, Y. Saito, S. Uchida, Y. Okada, A.C. Nairn, R.J. Davis, T. Yamamoto, M. Kinjo, H. Taru, and T. Suzuki. 2014. Quantitative analysis of APP axonal transport in neurons: Role of JIP1 in enhanced APP anterograde transport. Mol. Biol. Cell. 25:3569–3580. doi:10.1091/mbc.E14-06-1111.

Chiba, K., K.Y. Chien, Y. Sobu, S. Hata, S. Kato, T. Nakaya, Y. Okada, A.C. Nairn, M. Kinjo, H. Taru, R. Wang, and T. Suzuki. 2017. Phosphorylation of KLC1 modifies interaction with JIP1 and abolishes the enhanced fast velocity of APP transport by kinesin-1. Mol. Biol. Cell. 28:3857–3869. doi:10.1091/mbc.E17-05-0303.

Citron, M., T. Oltersdorf, C. Haass, L. McConlogue, A.Y. Hung, P. Seubert, C. Vigo-Pelfrey, I. Lieberburg, and D.J. Selkoe. 1992. Mutation of the β- amyloid precursor protein in familial Alzheimer’s disease increases β-protein production. Nature. 360:672–674. doi:10.1038/360672a0.

Citron, M., C. Vigo-Pelfrey, D.B. Teplow, C. Miller, D. Schenk, J. Johnston, B. Winblad, N. Venizelos, L. Lannfelt, and D.J. Selkoe. 1994. Excessive production of amyloid β-protein by peripheral cells of symptomatic and presymptomatic patients carrying the Swedish familial Alzheimer disease mutation. Proc. Natl. Acad. Sci. U. S. A. 91:11993–11997. doi:10.1073/pnas.91.25.11993.

Copenhaver, P.F., and J.M. Ramaker. 2016. Neuronal migration during development and the amyloid precursor protein. Curr. Opin. Insect Sci. 18:1– 10. doi:10.1016/j.cois.2016.08.001.

Das, U., D.A. Scott, A. Ganguly, E.H. Koo, Y. Tang, and S. Roy. 2013. Activity-induced convergence of app and bace-1 in acidic microdomains via an endocytosis-dependent pathway. Neuron. 79:447–460. doi:10.1016/j.neuron.2013.05.035.

Das, U., L. Wang, A. Ganguly, J.M. Saikia, S.L. Wagner, E.H. Koo, and S. Roy. 2015. Visualizing APP and BACE-1 approximation in neurons yields insight into the amyloidogenic pathway. Nat. Neurosci. 19:55–64. doi:10.1038/nn.4188.

El Meskini, R., L. Jin, R. Marx, A. Bruzzaniti, J. Lee, R.B. Emeson, and R.E. Mains. 2001. A signal sequence is sufficient for green Qluorescent protein to be routed to regulated secretory granules. Endocrinology. 142:864–873. doi:10.1210/endo.142.2.7929.

Ferreira, A., A. Caceres, and K.S. Kosik. 1993. Intraneuronal compartments of the amyloid precursor protein. J. Neurosci. 13:3112–3123. doi:10.1523/jneurosci.13-07-03112.1993.

Fourriere, L., and P.A. Gleeson. 2021. Amyloid β production along the neuronal secretory pathway: Dangerous liaisons in the Golgi? TrafCic. 1–9. doi:10.1111/tra.12808.

Frank, M., C.G. Citarella, G.B. Quinones, and M. Bentley. 2020. A Novel Labeling Strategy Reveals That Myosin Va and Myosin Vb Bind the Same Dendritically Polarized Vesicle Population. TrafCic. 21:689–701. doi:10.1111/tra.12764.

Frank, M., A.T. Nabb, S.P. Gilbert, and M. Bentley. 2022. Propofol attenuates kinesin-mediated axonal vesicle transport and fusion. Mol. Biol. Cell. 33:ar119. doi:10.1091/mbc.E22-07-0276.

Fu, M.M., and E.L.F. Holzbaur. 2013. JIP1 regulates the directionality of APP axonal transport by coordinating kinesin and dynein motors. J. Cell Biol. 202:495–508. doi:10.1083/jcb.201302078.

Ganguly, A., X. Han, U. Das, L. Wang, J. Loi, J. Sun, D. Gitler, G. Caillol, C. Leterrier, J.R. Yates, and S. Roy. 2017. Hsc70 chaperone activity is required for the cytosolic slow axonal transport of synapsin. J. Cell Biol. 216:2059–2074. doi:10.1083/jcb.201604028.

Garbouchian, A., A. Montgomery, S.P. Gilbert, and M. Bentley. 2022. KAP is the neuronal organelle adaptor for Kinesin-2 KIF3AB and KIF3AC. Mol. Biol. Cell. 33. doi:10.1091/mbc.e22-08-0336.

Goldstein, L.S.B., and U. Das. 2021. The cellular machinery of post-endocytic APP trafficking in Alzheimer’s disease: A future target for therapeutic intervention? 177. Elsevier Inc. 109–122 pp.

Greenfield, J.P., J. Tsai, G.K. Gouras, B. Hai, G. Thinakaran, F. Checler, S.S. Sisodia, P. Greengard, and H. Xu. 1999. Endoplasmic reticulum and trans-Golgi network generate distinct populations of Alzheimer beta-amyloid peptides. Proc. Natl. Acad. Sci. U. S. A. 96:742–747. doi:10.1073/pnas.96.2.742.

Haass, C., C. Kaether, G. Thinakaran, and S. Sisodia. 2012. Trafficking and proteolytic processing of APP. Cold Spring Harb. Perspect. Med. 2:1–26. doi:10.1101/cshperspect.a006270.

Haass, C., E.H. Koo, A. Mellon, A.Y. Hung, and D.J. Selkoe. 1992. Targeting of cell-surface beta-amyloid precursor protein to lysosomes: alternative processing into amyloid-bearing fragments. Nature. 357:500–503. doi:10.1038/357500a0.

Haass, C., C.A. Lemere, A. Capell, M. Citron, P. Seubert, D. Schenk, L. Lannfelt, and D.J. Selkoe. 1995. The Swedish mutation causes early-onset Alzheimer’s disease by β-secretase cleavage within the secretory pathway. Nat. Med. 1:1291–1296. doi:10.1038/nm1295-1291.

Heidemann, S.R., S. Kaech, R.E. Buxbaum, and A. Matus. 1999. Direct observations of the mechanical behaviors of the cytoskeleton in living fibroblasts. J. Cell Biol. 145:109–122.

Hunter, S., and C. Brayne. 2018. Understanding the roles of mutations in the amyloid precursor protein in Alzheimer disease. Mol. Psychiatry. 23:81–93. doi:10.1038/mp.2017.218.

Icking, A., M. Amaddii, M. Ruonala, S. Höning, and R. Tikkanen. 2007. Polarized transport of Alzheimer amyloid precursor protein is mediated by adaptor protein complex AP1-1B. TrafCic. 8:285–296. doi:10.1111/j.1600-0854.2006.00526.x.

Januário, Y.C., J. Eden, L.S. de Oliveira, R. De Pace, L.A. Tavares, M.E. da Silva-Januário, V.B. Apolloni, E.L. Wilby, R. Altmeyer, P.V. Burgos, S.A.L. Corrêa, D.C. Gershlick, and L.L.P. daSilva. 2022. Clathrin adaptor AP-1–mediated Golgi export of amyloid precursor protein is crucial for the production of neurotoxic amyloid fragments. J. Biol. Chem. 298:102172. doi:10.1016/j.jbc.2022.102172.

Johnston, J.A., R.F. Cowburn, S. Norgren, B. Wiehager, N. Venizelos, B. Winblad, C. Vigo-Pelfrey, D. Schenk, L. Lannfelt, and C. O’Neill. 1994. Increased β-amyloid release and levels of amyloid precursor protein (APP) in fibroblast cell lines from family members with the Swedish Alzheimer’s disease APP670/671 mutation. FEBS Lett. 354:274–278. doi:10.1016/0014-5793(94)01137-0.

Kaech, S., and G. Banker. 2006. Culturing hippocampal neurons. Nat. Protoc. 1:2406–2415. doi:10.1038/nprot.2006.356.

Kaech, S., C.F. Huang, and G. Banker. 2012a. General considerations for live imaging of developing hippocampal neurons in culture. Cold Spring Harb. Protoc. 7:312–318. doi:10.1101/pdb.ip068221.

Kaech, S., C.F. Huang, and G. Banker. 2012b. Short-term high-resolution imaging of developing hippocampal neurons in culture. Cold Spring Harb. Protoc. 7:340–343. doi:10.1101/pdb.prot068247.

Kaether, C., P. Skehel, and C.G. Dotti. 2000. Axonal membrane proteins are transported in distinct carriers: A two-color video microscopy study in cultured hippocampal neurons. Mol. Biol. Cell. 11:1213–1224. doi:10.1091/mbc.11.4.1213.

Kamal, A., G.B. Stokin, Z. Yang, C.H. Xia, and L.S.B. Goldstein. 2000. Axonal transport of amyloid precursor protein is mediated by direct binding to the kinesin light chain subunit of kinesin-I. Neuron. 28:449–459. doi:10.1016/S0896-6273(00)00124-0.

Kamiguchi, H., K.E. Long, M. Pendergast, A.W. Schaefer, I. Rapoport, T. Kirchhausen, and V. Lemmon. 1998. The neural cell adhesion molecule L1 interacts with the AP-2 adaptor and is endocytosed via the clathrin-mediated pathway. J. Neurosci. 18:5311–5321. doi:10.1523/jneurosci.18-14-05311.1998.

Kennedy, M.E., A.W. Stamford, X. Chen, K. Cox, J.N. Cumming, M.F. Dockendorf, M. Egan, L. Ereshefsky, R.A. Hodgson, L.A. Hyde, S. Jhee, H.J. Kleijn, R. Kuvelkar, W. Li, B.A. Mattson, H. Mei, J. Palcza, J.D. Scott, M. Tanen, M.D. Troyer, J.L. Tseng, J.A. Stone, E.M. Parker, and M.S. Forman. 2016. The BACE1 inhibitor verubecestat (MK-8931) reduces CNS β-amyloid in animal models and in Alzheimer’s disease patients. Sci. Transl. Med. 8:363ra150. doi:10.1126/scitranslmed.aad9704.

Koo, E.H., S.L. Squazzo, D.J. Selkoe, and C.H. Koo. 1996. Trafficking of cell-surface amyloid beta-protein precursor. I. Secretion, endocytosis and recycling as detected by labeled monoclonal antibody. J. Cell Sci. 109 (Pt 5):991–998. doi:10.1242/jcs.109.5.991.

Lai, A., S.S. Sisodia, and I.S. Trowbridge. 1995. Characterization of sorting signals in the beta-amyloid precursor protein cytoplasmic domain. J. iol. Chem. 270:3565–3573.

Lee, J., C. Retamal, L. Cuitiño A. Caruano-Yzermans, J.-E. Shin, P. van Kerkhof, M.-P. Marzolo, and G. Bu. 2008. Adaptor protein sorting nexin 17 regulates amyloid precursor protein trafficking and processing in the early endosomes. J. Biol. Chem. 283:11501–11508. doi:10.1074/jbc.M800642200.

Lorenzen, A., J. Samosh, K. Vandewark, P.H. Anborgh, C. Seah, A.C. Magalhaes, S.P. Cregan, S.S.G. Ferguson, and S.H. Pasternak. 2010. Rapid and Direct Transport of Cell Surface APP to the Lysosome defines a novel selective pathway. Mol. Brain. 3:1–13. doi:10.1186/1756-6606-3-11.

Matsuda, S., Y. Matsuda, and L. D’Adamio. 2003. Amyloid beta protein precursor (AbetaPP), but not AbetaPP-like protein 2, is bridged to the kinesin light chain by the scaffold protein JNK-interacting protein 1. J. Biol. Chem. 278:38601–38606. doi:10.1074/jbc.M304379200.

McCann, C.M., F.M. Bareyre, J.W. Lichtman, and J.R. Sanes. 2005. Peptide tags for labelling membrane proteins in live cells with multiple Qluorophores. Biotechniques. 38:945–952.

Montgomery, A.C., C.S. Mendoza, A. Garbouchian, G.B. Quinones, and M. Bentley. 2024. Polarized transport requires AP-1-mediated recruitment of KIF13A and KIF13B at the trans-Golgi. Mol. Biol. Cell. 35:ar61. doi:10.1091/mbc.E23-10-0401.

Morel, E., Z. Chamoun, Z.M. Lasiecka, R.B. Chan, R.L. Williamson, C. Vetanovetz, C. Dall’Armi, S. Simoes, K.S. Point Du Jour, B.D. McCabe, S.A. Small, and G. Di Paolo. 2013. Phosphatidylinositol-3-phosphate regulates sorting and processing of amyloid precursor protein through the endosomal system. Nat. Commun. 4:2250. doi:10.1038/ncomms3250.

Mullan, M., F. Crawford, K. Axelman, H. Houlden, L. Lilius, B. Winblad, and L. Lannfelt. 1992. Mullan, M., Crawford, F., Axelman, K., Houlden, H., Lilius, L., Winblad, B. & Lannfelt, L. (1992) A pathogenic mutation for probable Alzheimer’s disease in the APP gene at the N-terminus of beta-amyloid, Nat Genet. 1, 345-7. Nature. I:345–347.

Nabb, A.T., and M. Bentley. 2022. NgCAM and VAMP2 reveal that direct delivery and dendritic degradation maintain axonal polarity. Mol. Biol. Cell. 33:ar3. doi:10.1091/mbc.E21-08-0425.

Nabb, A.T., M. Frank, and M. Bentley. 2020. Smart motors and cargo steering drive kinesin-mediated selective transport. Mol. Cell. Neurosci. 103:103464. doi:10.1016/j.mcn.2019.103464.

Niederst, E.D., S.M. Reyna, and L.S.B. Goldstein. 2015. Axonal amyloid precursor protein and its fragments undergo somatodendritic endocytosis and processing. Mol. Biol. Cell. 26:205–217. doi:10.1091/mbc.E14-06-1049.

Nielsen, M.S., C. Gustafsen, P. Madsen, J.R. Nyengaard, G. Hermey, O. Bakke, M. Mari, P. Schu, R. Pohlmann, A. Dennes, and C.M. Petersen. 2007. Sorting by the cytoplasmic domain of the amyloid precursor protein binding receptor SorLA. Mol. Cell. Biol. 27:6842–6851. doi:10.1128/MCB.00815-07.

Nordstedt, C., G.L. Caporaso, J. Thyberg, S.E. Gandy, and P. Greengard. 1993. Identification of the Alzheimer beta/A4 amyloid precursor protein in clathrin-coated vesicles purified from PC12 cells. J. Biol. Chem. 268:608–612.

Norstrom, E.M., C. Zhang, R. Tanzi, and S.S. Sisodia. 2010. Identification of NEEP21 as a β-amyloid precursor protein-interacting protein in vivo that modulates amyloidogenic processing in vitro. J. Neurosci. 30:15677–15685. doi:10.1523/JNEUROSCI.4464-10.2010.

O’Brien, R.J., and P.C. Wong. 2011. Amyloid precursor protein processing and Alzheimer’s disease. Annu. Rev. Neurosci. 34:185–204. doi:10.1146/annurev-neuro-061010-113613.

Perez, R.G., S.L. Squazzo, and E.H. Koo. 1996. Enhanced release of amyloid β-protein from codon 670/671 “Swedish” mutant β-amyloid precursor protein occurs in both secretory and endocytic pathways. J. Biol. Chem. 271:9100– 9107. doi:10.1074/jbc.271.15.9100.

Petersen, J.D., S. Kaech, and G. Banker. 2014. Selective microtubule-based transport of dendritic membrane proteins arises in concert with axon specification. J. Neurosci. Off. J. Soc. Neurosci. 34:4135–47. doi:10.1523/jneurosci.3779-13.2014.

Ramaker, J.M., R.S. Cargill, T.L. Swanson, H. Quirindongo, M. Cassar, D. Kretzschmar, and P.F. Copenhaver. 2016. Amyloid Precursor Proteins Are Dynamically Trafficked and Processed during Neuronal Development. Front. Mol. Neurosci. 9:130. doi:10.3389/fnmol.2016.00130.

Ramaker, J.M., T.L. Swanson, and P.F. Copenhaver. 2013. Amyloid precursor proteins interact with the heterotrimeric G protein Go in the control of neuronal migration. J. Neurosci. Off. J. Soc. Neurosci. 33:10165–10181. doi:10.1523/JNEUROSCI.1146-13.2013.

Sampo, B., S. Kaech, S. Kunz, and G. Banker. 2003. Two distinct mechanisms target membrane proteins to the axonal surface. Neuron. 37:611–24. doi:10.1016/s0896-6273(03)00058-8.

Schmidt, V., A. Sporbert, M. Rohe, T. Reimer, A. Rehm, O.M. Andersen, and T.E. Willnow. 2007. SorLA/LR11 regulates processing of amyloid precursor protein via interaction with adaptors GGA and PACS-1. J. Biol. Chem. 282:32956– 32964. doi:10.1074/jbc.M705073200.

Selkoe, D.J., and J. Hardy. 2016. The amyloid hypothesis of Alzheimer’s disease at 25 years. EMBO Mol. Med. 8:595–608. doi:10.15252/emmm.201606210.

Shin, J.-Y., S.-B. Yu, U.-Y. Yu, S.-M. Ahnjo, and J.-H. Ahn. 2010. Swedish mutation within amyloid precursor protein modulates global gene expression towards the pathogenesis of Alzheimer’s disease. BMB Rep. 43:704–709. doi:10.5483/bmbrep.2010.43.10.704.

Silverman, M.A., S. Kaech, E.M. Ramser, X. Lu, M.R. Lasarev, S. Nagalla, and G. Banker. 2010. Expression of kinesin superfamily genes in cultured hippocampal neurons. Cytoskeleton. 67:784–795. doi:10.1002/cm.20487.

Sun, J., J. Carlson-Stevermer, U. Das, M. Shen, M. Delenclos, A.M. Snead, S.Y. Koo, L. Wang, D. Qiao, J. Loi, A.J. Petersen, M. Stockton, A. Bhattacharyya, M.V. Jones, X. Zhao, P.J. McLean, A.A. Sproul, K. Saha, and S. Roy. 2019. CRISPR/Cas9 editing of APP C-terminus attenuates β-cleavage and promotes α-cleavage. Nat. Commun. 10:53. doi:10.1038/s41467-018-07971-8.

Sun, J., and S. Roy. 2018. The physical approximation of APP and BACE-1: A key event in alzheimer’s disease pathogenesis. Dev. Neurobiol. 78:340– 347. doi:10.1002/dneu.22556.

Szodorai, A., Y.H. Kuan, S. Hunzelmann, U. Engel, A. Sakane, T. Sasaki, Y. Takai, J. Kirsch, U. Müller, K. Beyreuther, S. Brady, G. Morfini, and S. Kins. 2009. APP anterograde transport requires Rab3A GTPase activity for assembly of the transport vesicle. J. Neurosci. 29:14534–14544. doi:10.1523/JNEUROSCI.1546-09.2009.

Tam, J.H.K., C. Seah, and S.H. Pasternak. 2014. The amyloid precursor protein is rapidly transported from the golgi apparatus to the lysosome and where it is processed into beta-Amyloid. Mol. Brain. 7:1–18. doi:10.1186/s13041-014-0054-1.

Tambini, M.D., K.A. Norris, and L. D’Adamio. 2020. Opposite changes in APP processing and human aβ levels in rats carrying either a protective or a pathogenic APP mutation. eLife. 9:1–25. doi:10.7554/eLife.52612.

Tan, J.Z.A., and P.A. Gleeson. 2019a. The role of membrane trafficking in the processing of amyloid precursor protein and production of amyloid peptides in Alzheimer’s disease. Biochim. Biophys. Acta - Biomembr. 1861:697– 712. doi:10.1016/j.bbamem.2018.11.013.

Tan, J.Z.A., and P.A. Gleeson. 2019b. The trans-Golgi network is a major site for alpha-secretase processing of amyloid precursor protein in primary neurons. J. Biol. Chem. 294:1618–1631. doi:10.1074/jbc.RA118.005222.

Tang, Y., D.A. Scott, U. Das, S.D. Edland, K. Radomski, E.H. Koo, and S. Roy. 2012. Early and Selective Impairments in Axonal Transport Kinetics of Synaptic Cargoes Induced by Soluble Amyloid β-Protein Oligomers. TrafCic. 13:681–693. doi:10.1111/j.1600-0854.2012.01340.x.

Taylor, C.A., B.R. Miller, S.S. Shah, and C.A. Parish. 2016. A molecular dynamics study of the binary complexes of APP, JIP1, and the cargo binding domain of KLC. Proteins Struct. Funct. Bioinforma. doi:10.1002/prot.25208.

Thinakaran, G., and E.H. Koo. 2008. Amyloid precursor protein trafficking, processing, and function. J. Biol. Chem. 283:29615–29619. doi:10.1074/jbc.R800019200.

Thinakaran, G., D.B. Teplow, R. Siman, B. Greenberg, and S.S. Sisodia. 1996. Metabolism of the “Swedish” amyloid precursor protein variant in neuro2a (N2a) cells. Evidence that cleavage at the “beta-secretase” site occurs in the golgi apparatus. J. Biol. Chem. 271:9390–9397. doi:10.1074/jbc.271.16.9390.

Toh, W.H., J.Z.A. Tan, K.L. ZulkeQli, F.J. Houghton, and P.A. Gleeson. 2017. Amyloid precursor protein traffics from the Golgi directly to early endosomes in an Arl5b- and AP4-dependent pathway. TrafCic. 18:159–175. doi:10.1111/tra.12465.

Vassar, R., B.D. Bennett, S. Babu-Khan, S. Kahn, E.A. Mendiaz, P. Denis, D.B. Teplow, S. Ross, P. Amarante, R. Loeloff, Y. Luo, S. Fisher, J. Fuller, S. Edenson, J. Lile, M.A. Jarosinski, A.L. Biere, E. Curran, T. Burgess, J.C. Louis, F. Collins, J. Treanor, G. Rogers, and M. Citron. 1999. β-Secretase cleavage of Alzheimer’s amyloid precursor protein by the transmembrane aspartic protease BACE. Science. 286:735–741. doi:10.1126/science.286.5440.735.

Verwey, N.A., J.J.M. Hoozemans, C. Korth, M.R. van Royen, I. Prikulis, D. Wouters, H.A.M. TwaalQhoven, E.S. van Haastert, D. Schenk, P. Scheltens, A.J.M. Rozemuller, M.A. Blankenstein, and R. Veerhuis. 2013. Immunohistochemical characterization of novel monoclonal antibodies against the N-terminus of amyloid β-peptide. Amyloid Int. J. Exp. Clin. Investig. Off. J. Int. Soc. Amyloidosis. 20:179–187. doi:10.3109/13506129.2013.797389.

Wang, J., P.A. Gleeson, and L. Fourriere. 2024. Spatial-Temporal Mapping Reveals the Golgi as the Major Processing Site for the Pathogenic Swedish APP Mutation: Familial APP Mutant Shifts the Major APP Processing Site. TrafCic Cph. Den. 25:e12932. doi:10.1111/tra.12932.

Watson, E.T., W.R. Wegeng, S. Aravani, A.M. Ernst, and J. von Blume. 2025. Mechanistic insights into cargo sorting and export from the Golgi apparatus. Nat. Rev. Mol. Cell Biol. doi:10.1038/s41580-025-00907-3.

Woodruff, G., S.M. Reyna, M. Dunlap, R. Van Der Kant, J.A. Callender, J.E. Young, E.A. Roberts, and L.S.B. Goldstein. 2016. Defective Transcytosis of APP and Lipoproteins in Human iPSC-Derived Neurons with Familial Alzheimer’s Disease Mutations. Cell Rep. 17:759–773. doi:10.1016/j.celrep.2016.09.034.

Xia, W., W.J. Ray, B.L. Ostaszewski, T. Rahmati, W.T. Kimberly, M.S. Wolfe, J. Zhang, A.M. Goate, and D.J. Selkoe. 2000. Presenilin complexes with the C-terminal fragments of amyloid precursor protein at the sites of amyloid beta-protein generation. Proc. Natl. Acad. Sci. U. S. A. 97:9299–9304. doi:10.1073/pnas.97.16.9299.

Xu, H., D. Sweeney, R. Wang, G. Thinakaran, A.C. Lo, S.S. Sisodia, P. Greengard, and S. Gandy. 1997. Generation of Alzheimer beta-amyloid protein in the trans-Golgi network in the apparent absence of vesicle formation. Proc. Natl. Acad. Sci. U. S. A. 94:3748–3752. doi:10.1073/pnas.94.8.3748.

Yamazaki, T., D.J. Selkoe, and E.H. Koo. 1995. Trafficking of cell surface β-amyloid precursor protein: Retrograde and transcytotic transport in cultured neurons. J. Cell Biol. 129:431–442. doi:10.1083/jcb.129.2.431.

Yang, R., Z. Bostick, A. Garbouchian, J. Luisi, G. Banker, and M. Bentley. 2019. A novel strategy to visualize vesicle-bound kinesins reveals the diversity of kinesin-mediated transport. TrafCic. 20:851–866. doi:10.1111/tra.12692.

Yap, C.C., R.L. Nokes, D. Wisco, E. Anderson, H. Folsch, and B. Winckler. 2008. Pathway selection to the axon depends on multiple targeting signals in NgCAM. J Cell Sci. 121:1514–1525. doi:10.1242/jcs.022442.

Yildiz, A. 2024. Mechanism and regulation of kinesin motors. Nat. Rev. Mol. Cell Biol. 52:291–315. doi:10.1038/s41580-024-00780-6.

